# Genetic Deletion of TrpV1 and TrpA1 Does not Alter Avoidance of or Patterns of Brainstem Activation to Citric Acid in Mice

**DOI:** 10.1101/842112

**Authors:** Tian Yu, Courtney E. Wilson, Jennifer M. Stratford, Thomas E Finger

## Abstract

Exposure of the oral cavity to acidic solutions evokes not only a sensation of sour, but also of sharp or tangy. Acidic substances potentially stimulate both taste buds and acid-sensitive mucosal free nerve endings. Mice lacking taste function (P2X2/P2X3 double-KO mice) refuse acidic solutions similarly to wildtype mice and intraoral infusion of acidic solutions in these KO animals evokes substantial c-Fos activity within orosensory trigeminal nuclei as well as of the nucleus of the solitary tract (nTS) (Stratford et. al 2017). This residual acid-evoked, non-taste activity includes areas that receive inputs from trigeminal and glossopharyngeal peptidergic (CGRP-containing) nerve fibers that express TrpA1 and TrpV1 both of which are activated by low pH. We compared avoidance responses in wildtype (WT) and TrpA1/V1 double KO (TRPA1/V1^Db1-/-^) mice in brief-access behavioral assay (lickometer) to 1, 3, 10, 30 mM citric acid, along with 100 μM SC45647 and H_2_O. Both WT and TRPA1/V1^Db1-/-^ show similar avoidance, including to higher concentrations of citric acid (10 and 30 mM; pH 2.62 and pH 2.36 respectively), indicating that neither TrpA1 nor TrpV1 is necessary for the acid avoidance behavior in animals with an intact taste system. Similarly, induction of c-Fos in the nTS and dorsomedial spinal trigeminal nucleus was similar in the WT and TRPA1/V1^Db1-/-^animals. Taken together these results suggest non-TrpV1 and non-TrpA1 receptors underlie the residual responses to acids in mice lacking taste function.

## INTRODUCTION

When acidic solutions (e.g. lemon juice) are taken into the mouth, we describe the resulting perception as “sour” and consider it to be an aversive basic taste. However, the sensations arising from acidic substances in the oral cavity are a compound sensation of sour (a taste originating from taste buds) and direct activation of acid-sensitive general mucosal afferents of the oropharynx including the trigeminal, glossopharyngeal and superior laryngeal nerves. Indeed, a dictionary definition of sour (thefreedictionary.com) includes descriptors such ‘sharp,’ ‘tart’, or ‘tangy’ which are not taste modalities, but rather chemesthetic ones corresponding to activation of acid-sensitive mucosal nerves likely including polymodal nociceptors. Thus, aversion to acidic solutions might be due to activation of these nociceptors rather than of the taste system alone.

In mice lacking a functional taste system (P2X2/3-dbl-KO mice), the chorda tympani nerve, which provides only intragemmal innervation to taste buds (Silverman and Kruger 1990, Ohman-Gault, Huang et al. 2017), shows no responses to any tastants including acids although tactile and thermal responses remain (Finger, Danilova et al. 2005). Consistent with this, these mice show no preference for sweeteners or avoidance of bitter substances in brief-access taste tests. Despite the apparent lack of taste responses to acids, the P2X2/3-dbl-KO mice do exhibit normal avoidance of citric acid in similar brief access tests (Hallock, Tatangelo et al. 2009). This avoidance may be mediated not by taste, which is non-functional in these mice, but by acid-responsive fibers in the trigeminal, glossopharyngeal or laryngeal nerves which do show residual low level activity in the P2X2/3-dbl-KO mice (Ohkuri, Horio et al. 2012).

Polymodal nociceptors respond to acidification as well as other potentially painful stimuli (Bessou and Perl 1969). Many small caliber polymodal nociceptors that innervate the oral cavity (Kichko, Neuhuber et al. 2018, Wu, Arris et al. 2018) express one or both of the pH-sensitive transient receptor potential (Trp) channels, TrpA1 (Wang, Chang et al. 2011) and TrpV1 (Tominaga, Caterina et al. 1998). TrpA1 has been implicated especially in responsiveness to weak acids capable of penetrating cell membranes to produce intracellular acidification (Wang, Chang et al. 2011). TrpV1 is implicated in responses of the glossopharyngeal and vagus nerves to acidification of the oral and pharyngeal epithelium (Arai, Ohkuri et al. 2010). Gating of either of these channels by acids can directly activate the nociceptive fiber resulting in a noxious sensation suitable for driving avoidance behavior. Ablation of the ganglion cells expressing TrpV1 in mice lacking sour taste receptors does result in loss of acid-avoidance behavior (Zhang, Jin et al. 2019) but it is not clear whether this is due to loss of TrpV1 itself or other ion-sensing mechanisms of these neurons. We tested whether genetic deletion of both TrpA1 and TrpV1 affected either acid-induced avoidance behaviors or the pattern of neuronal activation in the brainstem taste (nuc. solitary tract, nTS) or trigeminal oral (dorsomedial trigeminal nucleus, DMSp5) sensory nuclei.

## Materials and Methods

### Animals

TrpA1V1-double KO mice, on a C57BL6 background (TRPA1/V1^Db1-/-^) were a generous gifts from Diana Bautista, University of California Berkeley (Gerhold and Bautista 2008). In these mice, deletion of TrpA1 had been accomplished by elimination of residues 901–951 including most of exon 23 – a region encoding the putative pore and part of the sixth transmembrane domain (Bautista, Jordt et al. 2006). The VR1 gene was disrupted (Caterina, Leffler et al. 2000) by deleting part of the fifth and all of the sixth putative transmembrane domains, together with the intervening pore-loop region. The TRPA1/V1^Db1-/-^ mice were genotyped by PCR to ensure quality control of the breeding. C57BL6/J mice were purchased from The Jackson Laboratory (Bar Harbor, ME). For brief access lickometer behavioral assays, 2-6 months old male (n=9) and female (n=6) TRPA1/V1^Db1-/-^ mice and 2-10 month old male (n=8) and female (n=3) C57BL6/J mice were used. For immunohistochemical staining, we used TRPA1/V1^Db1-/-^ (2 to 8 mo, n=10, 7 male and 3 female)) and C57BL6/J mice (4 to 7 mo, n=6, all male). In an addition to the C57 cases, we utilized 2 mice (1 male, 1 female) of the line B6.Cg-Tg(Fos-tTA,Fos-EGFP*)1Mmay/J which carry 2 transgenes associated with c-Fos: cfos-tTA and cfos-shEGFP. In these mice, on a C57BL6/J background, the randomly inserted transgenes utilize the c-Fos promotor to drive expression of respectively tTA and a short-lived (two-hour half-life) GFP. We did not find close correlation between GFP expression and c-Fos immunostaining and so the GFP results were ignored in our analysis. Nonethelss, the c-Fos counts from these animals were entirely consistent with those obtained from the wildtype C57BL6/J mice and so were included in those results.

All animal procedures were performed in accordance with NIH guidelines and were approved by the Institutional Animal Care and Use Committee at the University of Colorado School of Medicine.

### Brief access preference tests

Davis Rig Lickometers (MS-160; Dilog Instruments and Systems) were used for training and brief-access testing of mice. The animals underwent a 4-day water training before acid testing. Mice were placed on water deprivation 20 hours before the first day of training or acid testing. On the first day of the 4-day training, water-deprived animals were allowed access to a single water spout for a 30-min period. In the following three days of water training, 2-4 bottles of water were given in random sequences for 15 min. Mice had 5 seconds to lick at each spout after their first lick before the door closed and the bottle was switched to the next position with an inter-trial interval of 7.5 s. The mice were considered well-trained if they lick more than 30 times consistently during each trial for the first 15 trials. No mice lost more than 15% of body weight during any phase of the experimental procedure. All mice were given 2 day recovery period with ad libitum access to water after every 4 day block of testing.

For the acid testing days, six bottles of tastants including 4 concentrations of acetic or citric acid (1 mM, 3 mM, 10 mM and 30 mM), 1 concentration of artificial sweetener SC45647 (100 μM or 300μM) and H_2_O were presented in a testing block with a set up similar to the training session. Testing periods lasted for 15 min, with the opportunity for the mice to sample from the 6 solutions a total of 30 times. Licks were measured using InstaCal software. Each block of 6 tastant trials featured each solution in random order. Only data from completed blocks were used in the calculation of preference. Preference was calculated by averaging the licks per tastant block of each solution relative to those for water (i.e., a ‘lick ratio’). The same mice that were assessed for citric acid preference were then tested for acetic acid preference again with the same paradigm. The solutions used in acetic acid testing were as follows: 100 or 300 μM SC45647, H_2_O, 1, 3, 10 and 30 mM acetic acid. Testing for each series of tastants was repeated at least twice. Because there was no statistically significant difference in licking across each testing day for each mouse [F(1,2) = 0.42, p = 0.66], the number of licks for each tastant were averages across these testing days for each mouse.

### Citric acid stimulation for c-Fos via intraoral cannula

Both bilateral intraoral cannulae implantation and stimulation methods were adopted from (Stratford and Thompson 2016). Briefly, mice were anesthetized with an intramuscular injection of a combination of medetomidine hydrochloride (Domitor; 0.4 mg/kg; Pfizer) and ketamine hydrochloride (40 mg/kg; Bioniche Pharma). Intraoral cannulae were inserted via a midline incision immediately caudal to the pinnae. Then a sterile 2-3 cm stainless steel needle (19 gauge; Hamilton) with a flared end polyethylene tube (50 gauge; Becton Dickinson) was inserted from behind the pinna and guided subcutaneously into the oral cavity, lateral to the first maxillary molar. The needle was then withdrawn, leaving the polyethylene cannula and washer in place in the rear of the oral cavity. These mice were given 4 days for recovery prior to training.

For training and testing, liquids were delivered into intraoral cannula via a 5-cc syringe connected to a syringe pump (Model R99-E, Razel Scientific Instruments). To train the mice for the acclimation of liquid stimulation through the cannulae, they were water deprived for 23 hour / day, and given 3 ml of deionized water as a constant pulsatile flow of 0.1 ml/min through one of the two intraoral cannulae over the course of 30 min. We trained all the mice with deionized water for 2 days followed by stimulation with either deionized water (n=4 [all M] for C57BL6/J, n=5 [2 F, 3 M] for TrpA1V1-dbl KO), or 30 mM citric acid (n=4 [1 F, 3 M] for C57BL6/J, n=5 [1F, 4M] for TrpA1V1-dbl KO). Animals exposed to the stimulus were left undisturbed for an additional 45 min prior to sacrifice.

### c-Fos immunohistochemistry

Seventy-five minutes after onset of taste stimulation through cannula, animals were deeply anesthetized with Fatal-Plus^®^ (50 mg/kg intraperitoneally; MWI), and then perfused transcardially with saline followed by 4% paraformaldehyde (PFA) in 0.1 M pH 7.2 phosphate buffer (PB). The brains were post fixed for 3 hours at room temperature, and then cryo-protected overnight in 20% sucrose in PB at 4 °C. After cyroprotection, brainstems were cut and embedded in OCT, frozen and sectioned at 40 μm using a cryostat. Free-floating sections were collected in PBS.

For immunostaining, sections were washed in PBS 3 times and then processed for antigen retrieval in sodium citrate (pH 6) at 85 °C for 10 min. After tissues cooled, non-specific protein binding was blocked in a medium consisting of 2% normal donkey serum (Jackson ImmunoResearch) in antibody medium (AB medium: 0.3% TritonX100, 0.15 M NaCl, and 1% BSA in PB) for 1 hour at room temperature.

We utilized Mouse c-Fos antibody (1:1000, PhosphoSolutions; Cat#: 309-cFos; RRID RRID:AB_2632380; Lot#: GS117P) and Rabbit P2X2 antibody (1:1000, Alomone Labs; Cat#:APR003; RRID AB_2040054; Lot# APR003AN1002). These antibodies were diluted in AB medium accordingly and were used to incubate the sections for 4 days at 4 °C. After 3 10-min washes in PBS, sections were transferred to secondary antibody cocktail (Alexa Fluor 568 donkey anti mouse, 1:500; Alexa Fluor 488 donkey anti rabbit, 1:500; NeuroTrace Nissl 640/660, 1:500; all from Life Technologies) for 2-hour incubation at room temperature. Following another 3 10-min washes in PBS, free-floating tissue sections were mounted onto Microscope Slides (Tanner scientific, #TNR WHT90AD), and then coverslipped using Fuoromount-G (Southern Biotech).

According to the manufacturer’s data sheet, the PhosphoSolutions Mouse c-Fos antibody shows a broad band of reactivity in Western blots of HeLa cells centered at 50kDa. We tested whether this antibody stains similarly in fixed mouse brain tissues by comparing immunoreactivity of this antibody to the one we have utilized previously: we allowed a wildtype mouse to drink 150 mM MSG for 30min and perfused the mouse with 4% buffered paraformaldehyde after an additional 45 mins. The olfactory bulb & brainstem were removed, cryoprotected in 20% sucrose and frozen free-floating sections were cut at 40 μm. Representative sections were incubated in sodium citrate buffer pH6 at 85 °C for 10 min. After cooling to room temperature for 20 min, the sections were rinsed 3 × 5min in PBS, then incubated with 2%NDS+AB media for 1hr at room temperature. The free-floating sections then were incubated with a mixture of primary antibodies: rabbit a-cFos (lot.D00148958) 1:500 / mouse a-cFos (lot. GS418y) 1:1000 for 4 nights at 4 °C. After 3 × 10 min rinses in PBS, sections were incubated with a mixture of secondary antibodies: A488 donkey-a-rabbit 1:500/ A568 donkey-a-mouse 1:500 / Nissl 1:500 for 2 hrs at room temperature. The sections then were rinsed in PBS followed by 0.05M PB prior to coverslipping. The distribution of labeled cells within and around the nTS was similar to that observed in single label cases and nearly all cells were labeled by both antibodies. To quantitatively analyze the degree of co-localization, we counted cells labeled by one or both antibodies in the olfactory bulb where the density of labeled cells permits quantitative assessment. In randomly selected fields through the granule cells layer of 3 sections, we counted the number of single and double-label cells. Of 153 labeled cells, 149 were double-labeled; 4 were labeled by only the PhosphoSolutions antiserum and none were labeled only by the rabbit antibody. We conclude that the 2 antibodies label nearly identical populations and that results from the PhosphoSolutions antibody should be comparable to those obtained previously with the rabbit antibody.

### c-Fos activity determination

Brainstem sections were observed under an Olympus BX41microscope. Representative levels (r1, r2, r3, r4, i1, i2) of the nucleus of the solitary tract (nTS) were chosen under a 10X magnification objective according to Stratford et al. (Stratford, Thompson et al. 2017) and photographed at 20X magnification using CellSense software with a XM10 camera. Boundaries of the nTS were drawn using P2X2 and Nissl staining as reference and according to Stratford et al. (2017) in ImageJ. Boundaries of the DMSp5 were not distinct based on the images but the approximate boundaries were drawn using nTS boundaries and other brainstem nuclei as reference (Corson, Aldridge et al. 2012). The DMSp5 was taken as the area between the lateral edge of nTS and the spinal trigeminal nucleus.

The area of nTS was further divided into 6 parts following the system described in Stratford & Finger (Stratford and Finger 2011) using a horizontal line dividing the area into dorsal and ventral tiers and two vertical lines dividing the medial-lateral extent into thirds. The six parts of nTS were named DM, DI, DL, VM, VI, VL, and the cFos signals in these areas as well as DMSp5 were counted using ImageJ Cell counter plugin. The identity, stimulus and genotype of all cases were blinded to the person performing the cell counts.

### Statistical Analysis

Behavioral data are presented as group means with individual data points indicated. Immunohistochemical data are presented as group means ± SD. Data were first analyzed for normality using an Anderson-Darling test (AD). Because all data were deemed normally distributed based on the results of the AD test, appropriate two- and three-way analysis of variance (ANOVA)s were conducted (Statistica; StatSoft, Tulsa, OK). Tukey’s honest significant difference tests were used to assess statistically significant (p < 0.05) main effects or interactions (see Results for details).

## RESULTS

### Behavioral Assessment

For the acetic acid experiments, behavioral preferences were not statistically different between males and females across each genotype (WT and TRPA1/V1^Db1-/-^), (genotype x sex interaction, *F*(1, 14) = 0.00, *p* = 0.99). However, behavioral preferences were statistically different between different taste solutions, (main effect of taste, *F*(4, 56) = 24.89, *p* < 0.0001, Fig. 1A).

For the citric acid experiments, behavioral preferences were not statistically different between males and females, *F*(1, 22) = 2.80. *p* = 0.11. Moreover, the overall preference curves for both WT and TRPA1/V1^Db1-/-^ animals appeared nearly identical, (taste x genotype interaction, *F*(4,88) = 0.34, *p* = 0.85, Fig. 1B).

**Fig. 1:**
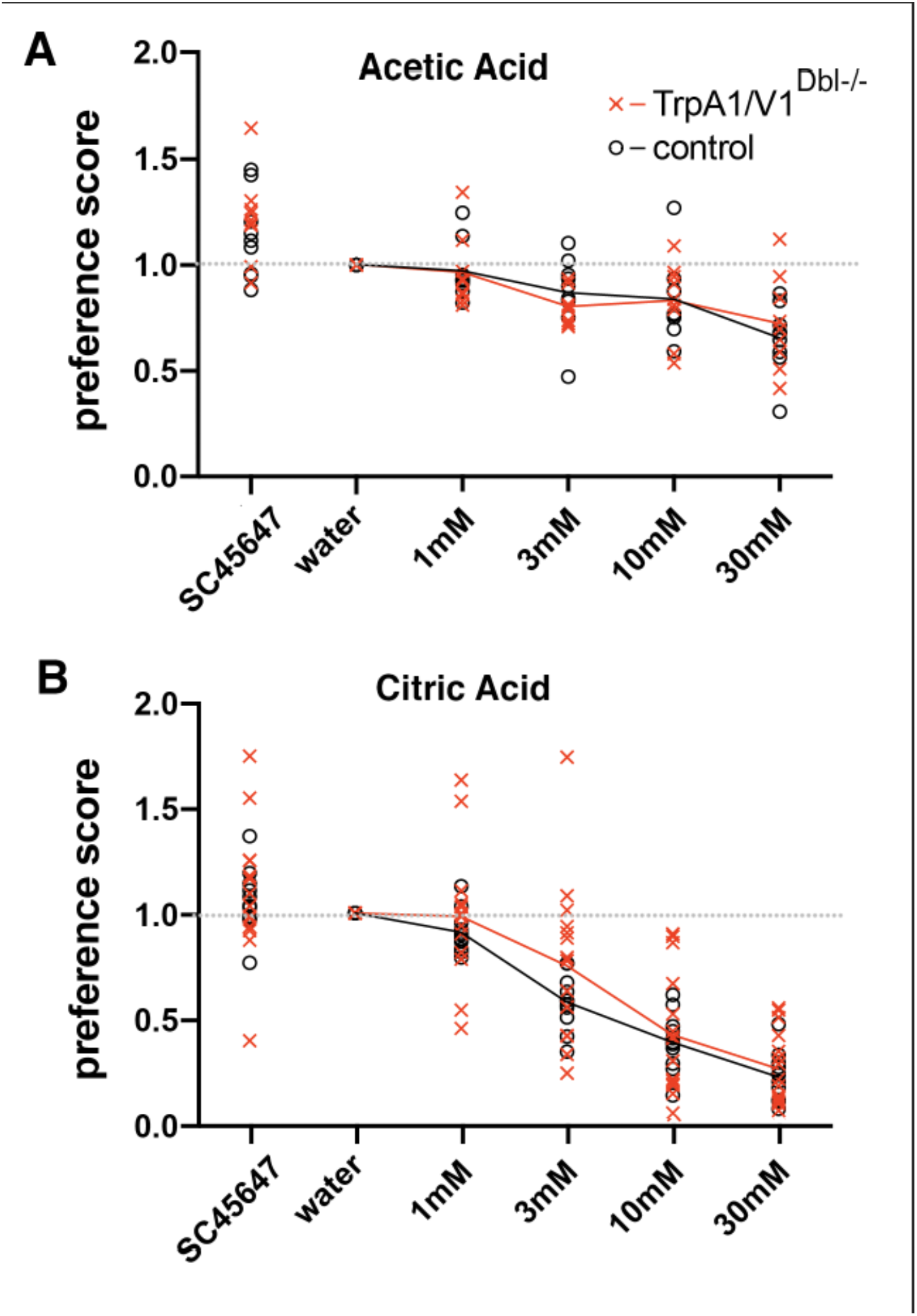
Brief access tests showing similar avoidance of acetic acid and citric acid for TRPA1/V1^Db1-/-^ mice and wildtype controls of the same background. ANOVA shows no significant difference between the groups.

### c-Fos Induction

Injection of 30 mM citric acid in the mouth via intraoral cannulae reliably induced c-Fos within particular regions of the nTS as compared to similar injection of water. As calculated previously, raw counts of c-Fos positive neurons in sub-compartments of the nTS were compared between water-stimulated and citric acid-stimulated animals to produce a measure of citric acid-specific c-Fos induction. In our previous study (Stratford, Thompson et al. 2017), acid-induced c-Fos was highest in the central portion of rostral-intermediate nTS (r3-i2) and in ventrolateral subdivisions in intermediate nTS (i1-i3). These regions did not show such activation in the P2X2/P2X3 dbl KO mice which lack taste function. However, in P2X2/P2X3 dbl KO animals, significant levels of activation to citric acid did remain in the DMSp5, which receives substantial input from polymodal nociceptor (CGRP+) fibers of the trigeminal and glossopharyngeal nerves. Accordingly, we especially focused on possible changes in c-Fos activation in this trigeminal-recipient area. Fig. 2 shows a representative image of c-Fos activation in relation to the lateral nTS and the DMSp5 of TRPA1/V1^Dbl-/-^ mice following stimulation with citric acid.

**Fig. 2:**
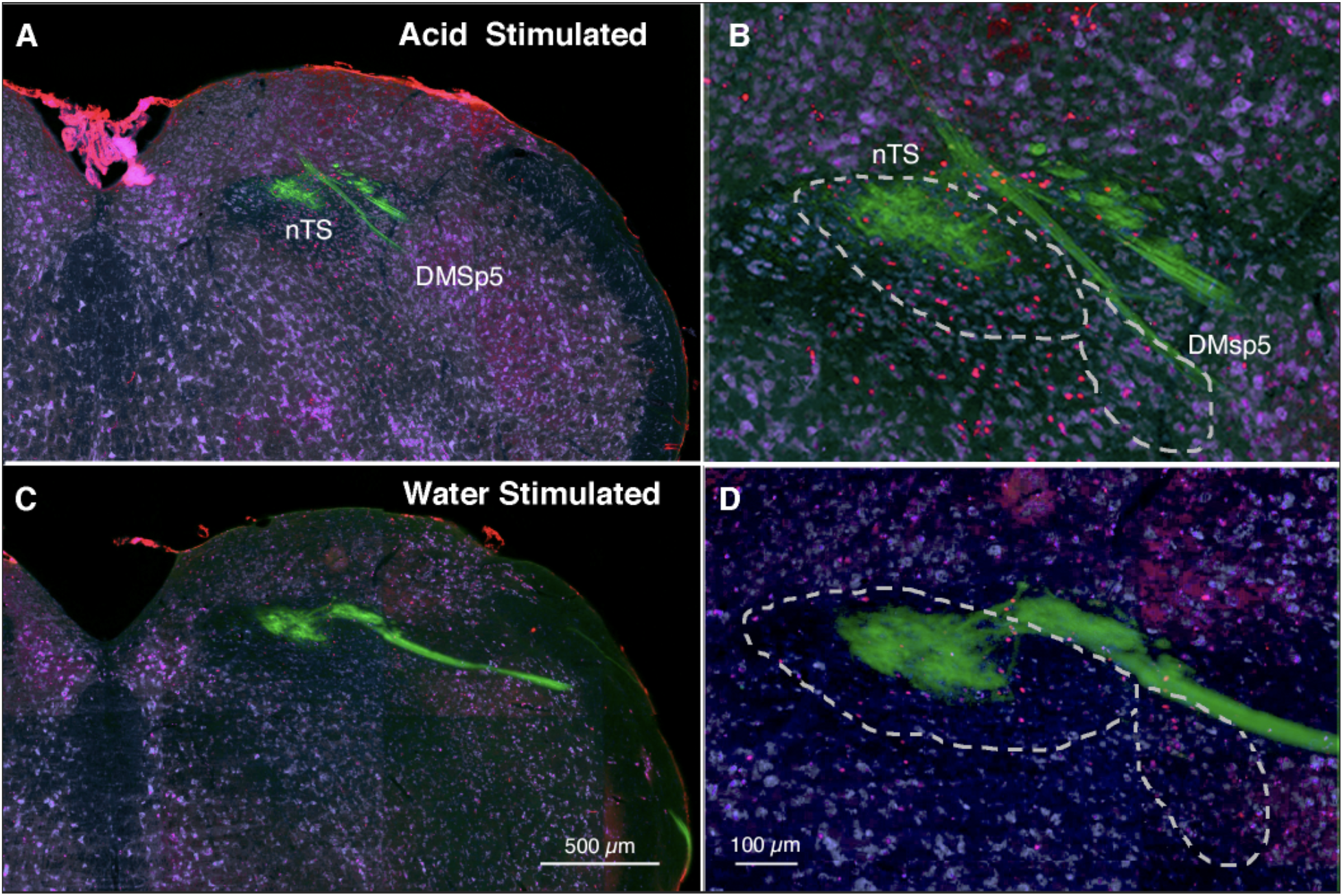
C-Fos staining (red dots) in nTS and DMsp5 of the TRPA1/V1^Db1-/-^ mouse at approximately the R3 level comparing a mouse stimulated by 30 mM citric acid (A, B) to one stimulated only by water (C, D). The number and location of c-fos positive cells is similar in WT and TRPA1/V1^Db1-/-^ strains. Note the higher number of c-fos positive cell nuclei (red) in the acid stimulated compared to the water-stimulated animal. Green staining shows the distribution of P2X3-immunoreactive fibers which terminate largely in the dorsocentral part of the nTS. Most acid-stimulated cells lie in the ventral tier of the nucleus and in the DMSp5 region ventrolateral to the nTS. Blue counterstain from FluoroNissl.

#### ‘Raw’ Tastant-Evoked c-Fos Counts

For all animals (i.e. both WT and TRPA1/V1^Dbl-^ mice), the total number of c-Fos positive cells evoked by citric acid oral stimulation was significantly greater than water stimulation within the nTS overall (water: 39.48 ± 3.59; citric acid: 71.03 ± 5.65, main effect of tastant: F (1, 5) = 49.53, p < 0.001; see Table 1). Moreover, the total number of water-evoked and citric acid-evoked c-Fos positive cells were significantly higher in WT as compared to TRPA1/V1^Db1-/-^ mice, regardless of tastant (For WT: 73.50 ± 7.17; For TRPA1/V1^Db1-/-^: 58.25 ± 5.57; main effect of genotype: F (1, 5) = 6.67, p < 0.05. However, there was no genotype x tastant interaction: *F*(1, 82) = 0.05, p = 0.83, see Table 1 and Fig. 3).

**Fig. 3.**
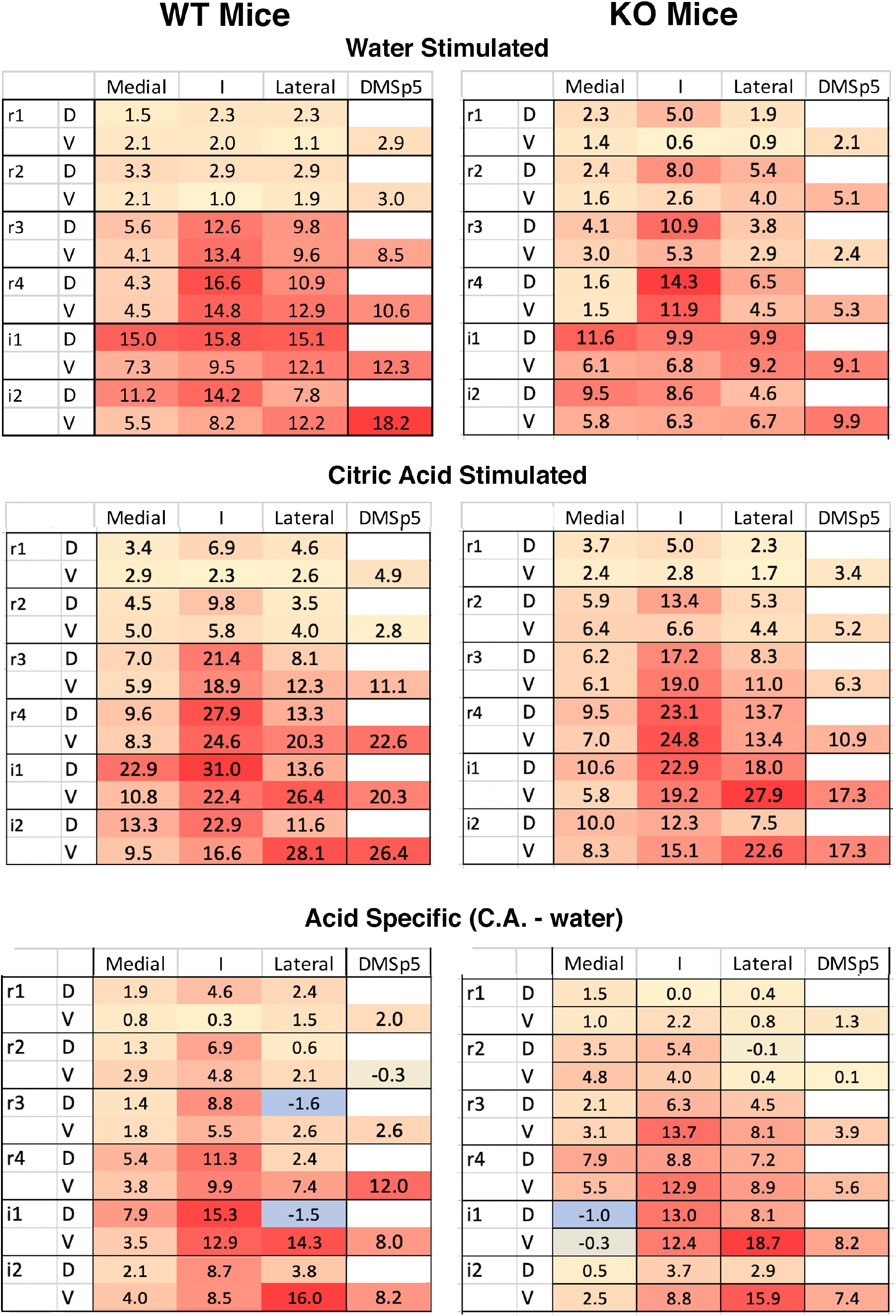
Heat maps of c-Fos expression in nTS and DMSp5 regions comparing wildtype (left) and KO (TRPA1/V1^Db1-/-^) mice, stimulated by intraoral injection of water (top row and by citric acid (middle row). The numbers indicate the aaverage number of c-fos-positive neurons in that compartment. See Suppl. Table 1 for means and standard deviations for each location for each genotype. Bottom row shows acid-specific activation, i.e. citric acid counts minus the counts produced by water alone. No significant differences occur either in the pattern or absolute level of expression across subregions (see text for details).

**Table 1.** Average (+/-S.D.) c-fos positive cells per region in the nTS and DMSp5 in WT and TrpA1/V1^Db1-/-^ mice following water and citric acid stimulation.

|  | Water | Citric Acid |
| --- | --- | --- |
| WT | 5.28 ± 1.58 | 9.83 ± 1.59 |
| TRPA1/V1<sup>Db1-/-</sup> | 13.72 ± 2.50 | 19.17 ± 2.54 |

Finally, there was no significant three-way tastant x genotype x nTS level interaction (*F* (1, 82) = 0.28, *p* = 0.92), nor was there a four-way tastant x genotype x nTS level x nTS subregion interaction (*F*(25, 410) = 0.55, *p* = 0.96).

#### Citric Acid-Specific c-Fos Activation

Because the number of c-Fos positive cells was lower in TRPA1/V1^Db1-/-^ mice as compared to WT nice (regardless of tastant), we calculated citric acid-specific activation for each region by taking the number of ‘raw’ citric acid-evoked c-Fos positive cells minus the average number of water-evoked c-Fos positive cells for each region for each genotype. Overall, citric-acid specific activation in the nTS did not differ between genotypes (main effect of genotype: *F* (1,5) = 0.08, *p* = 0.78). Furthermore, as shown in Fig. 3, the amount of citric acid-specific c-Fos was significantly different between nTS subregions and levels regardless of genotype, interaction between nTS subregion x nTS level: *F* (25, 210) = 4.24, p < 0.001. In particular, citric acid evoked significant activity in the central subregion of rostrointermediate nTS (r2-i2), in the lateral subregion of intermediate nTS (i1-2) and in the DMSp5 of both WT and TRPA1/V1^Db1-/-^ mice as compared to all other nTS subregions (all p’s < 0.05). This pattern is similar to that reported previously for the mixed background (Ola-C57/BL6) controls in the previous study (Stratford, Thompson et al. 2017).

In summary, the TRPA1/V1^Db1-/-^ mice show no differences in the pattern of neural activation by citric acid compared to the WT animals. This suggests that neither TrpA1 nor TrpV1 plays a significant role in detection and avoidance of 30 mM concentrations of citric acid, although this concentration is readily avoided by both TRPA1/V1^Dbl-/-^ and WT animals.

## DISCUSSION

Acidity is the underlying chemical feature of sour substances. When acids are taken into the mouth, they stimulate sour-sensitive taste cells which depolarize, then release neurotransmitter to activate the taste fibers innervating them. This signaling between taste cells and nerve fibers requires functional P2X2 and P2X3 receptors; genetic deletion or pharmacological blockage of these receptors essentially eliminates taste-mediated neural activity (Finger, Danilova et al. 2005, Vandenbeuch, Anderson et al. 2013). Similarly, taste driven acceptance of sweet and umami, and taste driven avoidance of bitter is lost in mice lacking functional P2X2 and P2X3 receptors. In contrast, behavioral avoidance of intraoral acids remains intact despite the loss of taste-related neural activity to these stimuli. Similarly, avoidance of acids persists even after genetic deletion of Otop1, the ion channel receptor for H^+^ underlying sour detection (Teng, Wilson et al. 2019, Zhang, Jin et al. 2019). These findings strongly suggest that not just taste, but another sensory modality drives the avoidance of acids in this context.

The oropharynx is innervated by chemically-sensitive free nerve endings arising from the trigeminal, glossopharyngeal and vagus nerves. These nerves include populations of polymodal nociceptors that express acid-sensitive ion channels including TrpA1 and TrpV1 (Tominaga, Caterina et al. 1998, Wang, Chang et al. 2011), and the TrpV1-expressing fibers are necessary for non-taste mediated avoidance of citric acid (Zhang, Jin et al. 2019). Yet these studies do not demonstrate that TrpV1 itself is the necessary receptor. It is likely that other acid-sensitive channels and receptors exist in these TrpV1-expressing fibers. Further, TrpV1 fully activates at a pH around 5 at body temperature (Tominaga, Caterina et al. 1998), whereas avoidance of citric acid begins at a pH near 3. It is likely that the tissue overlying the TrpV1 sensory terminals provides some buffering of acids applied to the surface of the epithelium, but weak acids, such as citric acid, effectively acidify the epithelium deep into the tissue (Richter, Caicedo et al. 2003), well beyond the region in which the nerve terminals lie. Taken together, these results suggest that TrpV1 itself may not be entirely responsible for responses to citric acid. Accordingly, we tested whether either TrpV1 or TrpA1 channels contribute to either the behavioral avoidance response, or the activation of brainstem neurons by citric acid. We found that neither the behavior nor the pattern and degree of neural activation was altered by genetic deletion of these channels.

If neither TrpV1 nor TrpA1 channels underlie avoidance of acidic substances in the absence of taste, what other mechanisms might be responsible? Likely candidates are one or more members of the ASIC (acid sensing ion channel) family, which are gated by decreases in extracellular pH. While many members of this family open at pH values near 7, others activate at lower pH values (Deval, Gasull et al. 2010). In particular, ASIC3 is expressed widely in polymodal nociceptors including those expressing TrpV1 (Trigeminal ganglion clusters 7 & 10 in (Nguyen, Wu et al. 2017)) and plays a role in responses to pH5.0 (Price, McIlwrath et al. 2001). Further, amiloride, a non-specific blocker of ASIC channels, decreases the irritation of citric acid (albeit at a much higher concentration) measured psychophysically (Dessirier, O’Mahony et al. 2000), supporting the role of ASICs in this response. Conversely, inclusion of amiloride, a blocker of ASICs, in orally-applied solutions in rats neither decreased the trigeminal response to citric acid nor attenuated acid-induced c-Fos in brainstem trigeminal nuclei (Sudo, Sudo et al. 2003). Whether ASICs or some other acid-responsive mechanisms play a role in non-gustatory behavioral avoidance of weak acids then is unresolved.

## ACKNOWLEDGEMENTS

The authors thank Mei Li for histological preparations and for comparison of antibody staining properties. We also thank NIDCD for supporting this investigation (RO1 DC012931).

---

**Supplemenatry Table 1:** Means and standard deviations of counts of c-fos positive cells in DMSp5 and each subnucleus of the NTS for water and citric acid stimulation by intraoral cannula for WT and TrpA1/V1^-/-^ mice.

| WT | H2O | DM | DI | DL | VM | VI | VL | DMSp5 |
| --- | --- | --- | --- | --- | --- | --- | --- | --- |
| **r1** |  | 1.5 | 2.3 | 2.3 | 2.1 | 2.0 | 1.1 | 2.9 |
| stdev |  | 1.1 | 1.8 | 1.0 | 0.8 | 1.1 | 1.1 | 2.7 |
| **r2** |  | 3.3 | 2.9 | 2.9 | 2.1 | 1.0 | 1.9 | 3.0 |
| stdev |  | 1.4 | 1.5 | 2.0 | 1.1 | 0.5 | 1.0 | 1.8 |
| **r3** |  | 5.6 | 12.6 | 9.8 | 4.1 | 13.4 | 9.6 | 8.5 |
| stdev |  | 4.3 | 6.3 | 5.9 | 1.9 | 7.6 | 6.0 | 9.5 |
| **r4** |  | 4.3 | 16.6 | 10.9 | 4.5 | 14.8 | 12.9 | 10.6 |
| stdev |  | 3.0 | 6.7 | 7.0 | 2.1 | 4.4 | 6.4 | 6.9 |
| **i1** |  | 15.0 | 15.8 | 15.1 | 7.3 | 9.5 | 12.1 | 12.3 |
| stdev |  | 6.2 | 8.7 | 6.3 | 5.3 | 4.5 | 6.8 | 8.2 |
| **i2** |  | 11.2 | 14.2 | 7.8 | 5.5 | 8.2 | 12.2 | 18.2 |
| stdev |  | 10.6 | 8.0 | 4.7 | 3.7 | 5.5 | 7.5 | 14.4 |

| WT | DM | DI | DL | VM | VI | VL | DMSp5 |
| --- | --- | --- | --- | --- | --- | --- | --- |
| **Cit Acid** |  |  |  |  |  |  |  |
| **r1** | 3.4 | 6.9 | 4.6 | 2.9 | 2.3 | 2.6 | 4.9 |
| stdev | 1.8 | 1.8 | 3.3 | 2.6 | 1.3 | 1.6 | 3.4 |
| **r2** | 4.5 | 9.8 | 3.5 | 5.0 | 5.8 | 4.0 | 2.8 |
| stdev | 1.6 | 3.4 | 2.0 | 2.7 | 4.3 | 2.1 | 2.9 |
| **r3** | 7.0 | 21.4 | 8.1 | 5.9 | 18.9 | 12.3 | 11.1 |
| stdev | 3.6 | 8.3 | 3.4 | 2.7 | 2.6 | 6.8 | 9.4 |
| **r4** | 9.6 | 27.9 | 13.3 | 8.3 | 24.6 | 20.3 | 22.6 |
| stdev | 3.4 | 9.4 | 3.3 | 3.9 | 6.0 | 7.4 | 14.3 |
| **i1** | 22.9 | 31.0 | 13.6 | 10.8 | 22.4 | 26.4 | 20.3 |
| stdev | 10.7 | 9.6 | 7.3 | 4.8 | 4.9 | 7.8 | 8.5 |
| **i2** | 13.3 | 22.9 | 11.6 | 9.5 | 16.6 | 28.1 | 26.4 |
| stdev | 9.0 | 4.8 | 5.4 | 5.4 | 7.5 | 9.8 | 18.8 |

| TrpA1/V1<sup>-/-</sup> |  | H2O | DM | DI | DL | VM | VI | VL | DMSp5 |
| --- | --- | --- | --- | --- | --- | --- | --- | --- | --- |
| **r1** |  |  | 2.3 | 5.0 | 1.9 | 1.4 | 0.6 | 0.9 | 2.1 |
|  | stdev |  | 3.4 | 4.6 | 2.0 | 1.5 | 0.7 | 1.7 | 2.9 |
| **r2** |  |  | 2.4 | 8.0 | 5.4 | 1.6 | 2.6 | 4.0 | 5.1 |
|  | stdev |  | 1.6 | 5.0 | 5.8 | 1.5 | 4.3 | 5.8 | 7.8 |
| **r3** |  |  | 4.1 | 10.9 | 3.8 | 3.0 | 5.3 | 2.9 | 2.4 |
|  | stdev |  | 2.3 | 4.8 | 1.9 | 2.7 | 5.2 | 2.7 | 2.1 |
| **r4** |  |  | 1.6 | 14.3 | 6.5 | 1.5 | 11.9 | 4.5 | 5.3 |
|  | stdev |  | 1.8 | 8.9 | 4.1 | 2.2 | 6.8 | 3.7 | 8.2 |
| **i1** |  |  | 11.6 | 9.9 | 9.9 | 6.1 | 6.8 | 9.2 | 9.1 |
|  | stdev |  | 6.1 | 5.9 | 8.0 | 3.4 | 5.2 | 7.2 | 11.6 |
| **i2** |  |  | 9.5 | 8.6 | 4.6 | 5.8 | 6.3 | 6.7 | 9.9 |
|  | stdev |  | 9.1 | 5.4 | 3.3 | 5.3 | 3.4 | 5.3 | 12.9 |

| TrpA1/V1<sup>-/-</sup> |  | Cit Acid | DM | DI | DL | VM | VI | VL | DMSp5 |
| --- | --- | --- | --- | --- | --- | --- | --- | --- | --- |
| **r1** |  |  | 3.7 | 5.0 | 2.3 | 2.4 | 2.8 | 1.7 | 3.4 |
|  | stdev |  | 2.8 | 2.0 | 1.1 | 0.8 | 2.4 | 1.7 | 2.2 |
| **r2** |  |  | 5.9 | 13.4 | 5.3 | 6.4 | 6.6 | 4.4 | 5.2 |
|  | stdev |  | 3.2 | 5.8 | 3.1 | 5.6 | 4.2 | 2.9 | 5.2 |
| **r3** |  |  | 6.2 | 17.2 | 8.3 | 6.1 | 19.0 | 11.0 | 6.3 |
|  | stdev |  | 3.9 | 8.0 | 5.3 | 4.4 | 9.2 | 8.7 | 4.7 |
| **r4** |  |  | 9.5 | 23.1 | 13.7 | 7.0 | 24.8 | 13.4 | 10.9 |
|  | stdev |  | 7.6 | 8.4 | 7.7 | 2.7 | 7.8 | 6.2 | 5.5 |
| **i1** |  |  | 10.6 | 22.9 | 18.0 | 5.8 | 19.2 | 27.9 | 17.3 |
|  | stdev |  | 5.3 | 18.3 | 9.2 | 3.2 | 9.8 | 12.0 | 10.9 |
| **i2** |  |  | 10.0 | 12.3 | 7.5 | 8.3 | 15.1 | 22.6 | 17.3 |
|  | stdev |  | 5.1 | 17.4 | 8.6 | 3.9 | 6.9 | 8.9 | 13.1 |

